# Serotonin reuptake inhibition rapidly enhances affect-reward coupling and improves mood over time

**DOI:** 10.64898/2026.07.03.736315

**Authors:** Regan Mills, Anais Durand, Meggy Croal, Robb B. Rutledge, Liam Mason

**Affiliations:** Research Department of Clinical, Educational and Health Psychology, University College London, London, UK; Department of Psychology, Yale University. New Haven, CT, USA

## Abstract

Selective serotonin reuptake inhibitors (SSRIs) are widely prescribed antidepressants, yet the mechanisms linking their rapid neurobiological effects to more gradual improvements in mood remain unclear. Computational accounts in which mood reflects the integration of reward outcome and reward prediction errors (RPEs) offer a compelling framework for examining SSRI effects, but reliance on laboratory tasks poses a major translational gap, both in timescale and ecological validity. We address this gap by examining the impact of SSRIs on affect-reward dynamics over a clinically relevant timescale using real-world rewards. Across 11 days, 66 healthy participants (n=37 receiving the SSRI citalopram) completed repeated ecological momentary assessments (EMA) of mood and goal-directed activities (up to 8 per day, totaling 3,542 surveys). By probing anticipated reward (pre-activity) and consummatory reward (post-activity), we derived real-world RPEs to test whether SSRIs amplify RPE impacts on momentary mood. Mixed-effects modelling showed that citalopram rapidly amplified the impact of reward outcome and RPEs on momentary mood, and predicted later increases in positive mood. Citalopram also selectively attenuated the impact of negative mood on expectations for future reward. These findings suggest that SSRIs modulate the bidirectional coupling between mood and reward processes in daily life, consistent with a positive feedback loop that could underpin the gradual mood improvement observed for antidepressant treatment. More broadly, our results highlight the utility of EMA as a highly sensitive real-world assay of early pharmacological effects on mood-reward dynamics.

## Introduction

Major depressive disorder is among the leading global causes of disability and disease burden, posing substantial clinical and economic challenges to public healthcare systems worldwide.^1^ Alongside psychological intervention, pharmacotherapy is a widely used treatment strategy for depression, with selective serotonin reuptake inhibitors (SSRIs) serving as a first-line antidepressant.^2^ While the neurochemical actions of SSRIs are well-established, the cognitive and computational mechanisms that translate their rapid monoaminergic effects into delayed clinical improvement remain unclear.^3,4^

Establishing the acute role of serotonin in affective processes, prior studies have typically used either a single acute dose or week-long treatment in healthy human volunteers.^5,6^ Specifically, short-term SSRI administration has been shown to produce a positive shift in affective processing at both behavioral and neural levels, by altering neural responses to emotional cues to elicit a “rosier” perspective.^7^ Such findings have laid the foundation for recent computational approaches of reinforcement-learning (RL), which can formalize how changes in affect-learning dynamics may contribute to gradual mood improvement during SSRI treatment.

In one such computational framework, Eldar et al. propose that mood is not simply reactive to external events but reflects an integration of recent reward prediction errors (RPEs, the discrepancy between expected and received outcomes).^8^ In turn, mood may act as a generalization mechanism, biasing the perception of subsequent outcomes to facilitate efficient learning in uncertain environments. Multiple laboratory-based studies have demonstrated this principle, showing that positive RPEs from unexpected financial rewards increase positive affect, which in turn elevates the subjective valuation of subsequent monetary rewards.^9,10^ In a seminal paper, Michely et al. extended these findings to suggest a mechanistic account of serotonin’s impact on mood, showing that citalopram increases the impact of experimentally induced positive affect on learning of novel, and reconsolidation of previously learned, reward associations.^11^ Therefore, rather than acting on affect or reward sensitivity directly, SSRIs amplify an interaction between the two.

However, it is not currently clear whether the proposed bidirectional relationship between affect and reward translates to the real world. Bridging this gap is essential for establishing the potential clinical relevance of this theoretical framework. In natural environments, reward operates across multiple dimensions, engaging partially distinct neural circuitry that support the processing of diverse social and non-social experiences.^12,13^ Mood is similarly dynamic and multifaceted, evolving over time through interactions between positive and negative affective states.^14^ Mood dynamics thus cannot be adequately captured by the single item ratings typically used in laboratory tasks (e.g., “How happy are you now?”). While such paradigms have provided important mechanistic insight into the interplay between affective and reward processes during experimental paradigms, they are limited in their ability to capture the forms of reward most relevant to real-world mood, and ultimately to the mechanisms central to SSRI antidepressant effects.

Ecological momentary assessment (EMA) provides a powerful approach for examining the dynamic impact of rewarding activities on mood, particularly as it enables real-time and multidimensional assessment of both constructs. Using EMA, Villano et al. demonstrated that outcomes and RPEs are key predictors of affect, using a single-shot emotionally salient event.^15^ Specifically, this study found that exam grade outcomes and prediction errors (PEs) about exam grades impacted the time course of both positive and negative affect, with the effect of these PEs persisting on the order of hours. They extended the same EMA paradigm to clinical contexts, investigating the impact of depression on affective responding to PEs, as well as the link between PE-based learning and negative emotionality.^16,17^ Moving forwards, it will be important to interrogate whether these mood-reward interactions extend beyond this specific experimental context to everyday goal-directed activities that are much more varied than exam grades.

In the present study, we investigated the impact of eight days of SSRI administration on momentary mood and reward dynamics. Leveraging the insights from well established computational models of mood, we measured pairs of reward expectations and outcomes over multiple daily activities to derive real-world RPEs, thus enabling us to examine the nature of mood dynamics using high-frequency and ecologically valid measures of goal-directed behaviour.^9,10^ Crucially, this approach also allowed us to determine whether EMA, informed by reinforcement learning models, can detect the early effects of SSRIs on the coupling between reward perception and momentary affect, therefore providing a sensitive and scalable readout of serotonergic effects that precedes and predict gradual symptom change.^18^

First, we sought to establish whether reward prediction errors from daily goal-directed activities would drive changes in momentary mood, extending prior laboratory and single-shot EMA findings to a fully naturalistic repeated reward context.^9,10,15–17^ Specifically, we predicted that activity-locked RPEs would drive independent changes in momentary mood over and above consummatory reward from the activity. Secondly, we predicted that SSRIs would boost the impact of these rewards on momentary mood. Leveraging the multi-day study design, we tested whether this effect on mood would strengthen over time. We also tested whether this effect would be stronger for positive rather than negative RPEs, consistent with prior evidence for valence-specific serotonergic effects.^11,19^ Lastly, in keeping with the idea that mood serves an adaptive role in biasing expectations about reward, we hypothesized that SSRI receipt would increase the coupling between momentary mood and reward expectation.^8,18^

## Results

### EMA-based mood and reward tracking across SSRI and placebo groups

70 healthy participants were recruited for this double-blind, placebo-controlled study, matched by age and gender across drug groups (mean age: 24.8 ± 5.9; range 18-39 years; 48 females; Table 1). 40 participants were randomly allocated to receive a daily oral dose of the SSRI citalopram (20 mg) over eight consecutive days, while the remaining 30 participants were allocated to placebo (Figure 1A). Participants also completed eleven days of EMA using the ‘The Happiness Project’ smartphone app (https://rutledgelab.org/), installed on their mobile phones during the in-person session on Day 1. They were expected to complete four pre-activity surveys per day, which surveyed momentary mood and a rewarding event assessment (Figure 1B). Importantly, each of these surveys generated a paired post-activity survey, yielding up to eight daily probes. Factor analysis was used to derive momentary positive and negative affect from pre- and post-activity mood surveys. RPEs were calculated from the difference between post-activity reward outcomes (r) and pre-activity reward predictions (ν) (see Methods for more detail).

**Figure 1.**
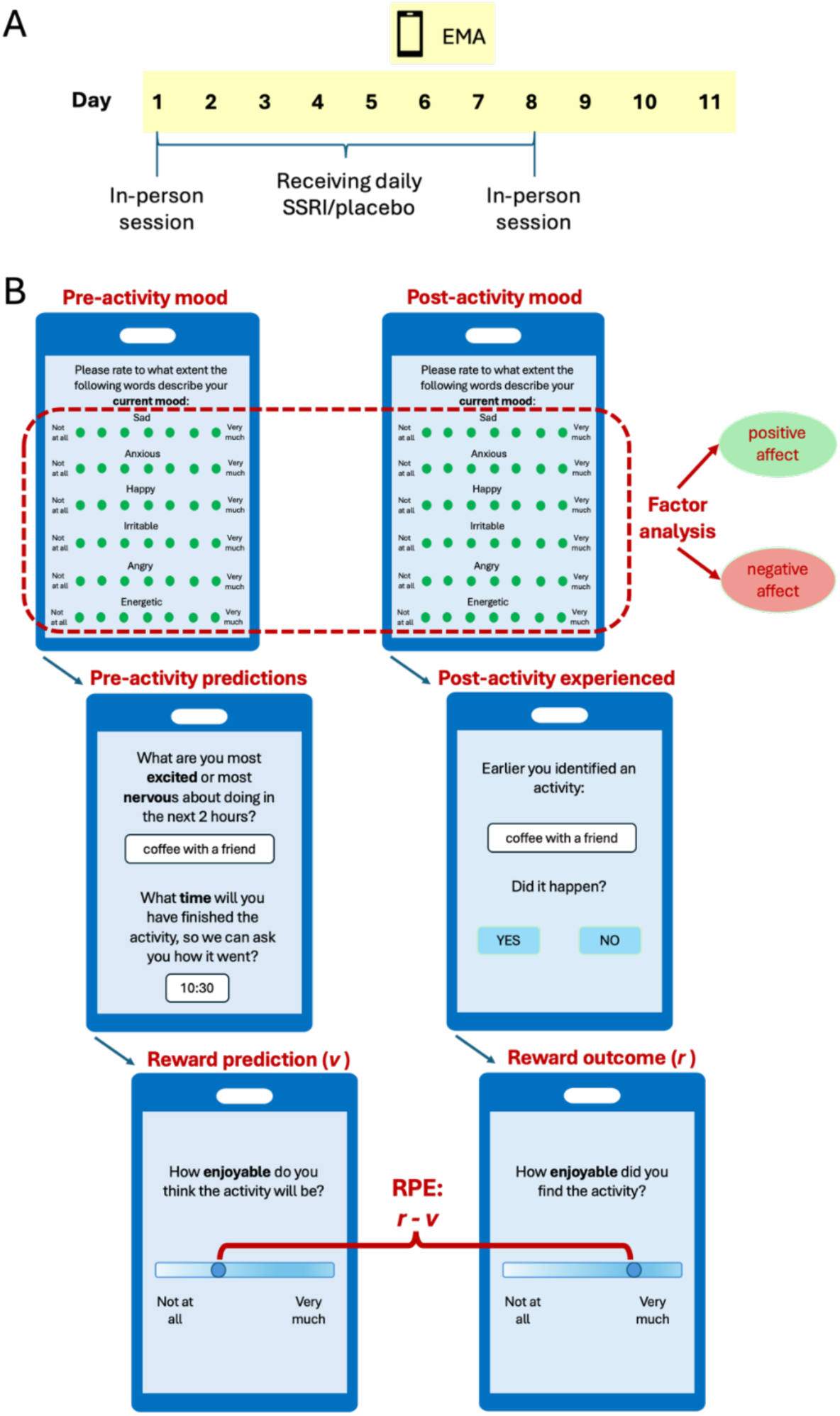
Pharmacological procedure and EMA design. **(A) Pharmacological procedure**, where participants were randomly assigned to receive either citalopram or placebo from Days 1 to 8. In-person study sessions were completed on the first and last day of pharmacological manipulation, allowing for the completion of mood and psychiatric symptom questionnaires and baseline testing. EMA was conducted from Days 1 to 11. **(B) EMA design**, showing example screenshots of the pre- and post-activity surveys as they appeared on participant mobile phones. Daily notifications for the pre-activity survey were scheduled at 9:00am, 12:00pm, 3:00pm and 6:00pm, and surveyed momentary mood and a rewarding event assessment. For this, participants were asked to identify a salient activity that would occur in the next two hours Representative examples of participant activities included “walking the dog”, and “getting lunch with a friend”. To trigger the paired post-activity survey, participants were then asked to identify the time that the activity would be finished, as well as rate the activity’s predicted reward. The post-activity survey was notified at the time specified by the participant, and assessed momentary mood, followed by probes for the actual experience of reward during the activity.

**Table 1.** Participant demographics and affective state questionnaire data.

|  | Full sample (n=70) |  |  | Final analysis sample (n=66) |  |  |
| --- | --- | --- | --- | --- | --- | --- |
|  | Placebo<br>(n=30) | SSRI<br>(n=40) | p-value | Placebo<br>(n=29) | SSRI<br>(n=37) | p-value |
| **Gender** (female), n (%) | 24 (80%) | 24 (60%) | 0.128 | 23 (79%) | 22 (59%) | 0.146 |
| **Age** (years), mean (SD) | 24.8 (5.9) | 24.8 (5.9) | 0.999 | 25.0 (6.1) | 25.0 (6.0) | 1.00 |
| **Weight** (kg), mean (SD) | 63.6 (12.9) | 66.2 (13.6) | 0.413 | 63.6 (13.2) | 67.3 (13.0) | 0.266 |
| **PHQ-9** (day 1), mean (SD) | 4.14 (3.1)<sup>a</sup> | 4.52 (4.5) | 0.692 | 3.93 (3.1)<sup>a</sup> | 4.24 (4.4) | 0.754 |
| **PHQ-9** (day 8), mean (SD) | 3.63 (2.3)<sup>b</sup> | 4.89 (4.6)<sup>c</sup> | 0.200 | 3.62 (2.4)<sup>b</sup> | 4.38 (4.1)<sup>c</sup> | 0.401 |
| **PHQ-9** (day 8 – day 1), mean (SD) | -0.37 (2.0)<sup>b</sup> | 0.44 (3.8)<sup>c</sup> | 0.310 | -0.31 (2.0)<sup>b</sup> | 0.26 (3.6)<sup>c</sup> | 0.464 |
| **GAD-7** (day 1), mean (SD) | 3.38 (2.8)<sup>a</sup> | 3.73 (3.6) | 0.666 | 3.19 (2.8)<sup>a</sup> | 3.29 (3.1) | 0.889 |
| **GAD-7** (day 8), mean (SD) | 2.89 (2.6)<sup>b</sup> | 3.06 (4.2)<sup>c</sup> | 0.855 | 2.92 (2.6)<sup>b</sup> | 2.65 (3.9)<sup>c</sup> | 0.757 |
| **GAD-7** (day 8 – day 1), mean (SD) | -0.41 (1.90)<sup>b</sup> | -0.67 (3.5)<sup>c</sup> | 0.617 | -0.31 (1.9)<sup>b</sup> | -0.59 (3.5)<sup>c</sup> | 0.603 |
| **AMI** (day 1), mean (SD) | 3.88 (1.1)<sup>a</sup> | 3.66 (1.1) | 0.414 | 3.78 (1.0)<sup>a</sup> | 3.62 (1.1) | 0.569 |
| **AMI** (day 8), mean (SD) | 4.08 (1.5)<sup>b</sup> | 3.75 (1.4)<sup>c</sup> | 0.363 | 3.97 (1.4)<sup>b</sup> | 3.71 (1.4)<sup>c</sup> | 0.481 |
| **AMI** (day 8 – day 1), mean (SD) | 0.25 (0.8)<sup>b</sup> | 0.25 (0.9)<sup>c</sup> | 0.992 | 0.19 (0.8)<sup>b</sup> | 0.26 (0.9)<sup>c</sup> | 0.634 |

**Note:** Missing participant data for: <sup>a</sup>n=1, <sup>b</sup>n=3, <sup>c</sup>n=4. PHQ-9 = Patient Health Questionnaire. GAD-7 = Generalized Anxiety Disorder scale. AMI = Apathy Motivation Index.

Participants who dropped out of the study after the in-person session on Day 1 were excluded from the dataset, leaving a final analysis sample of 66 participants. Survey response rates in the final analysis sample were comparable to EMA studies with similar sampling frequencies.^20^ Of the 2904 potential data points for pre-activity surveys (66 participants x 44 surveys), complete data were available for 2069 (71.2%). Of the potential post-activity surveys, 1473 (71.2%) were completed. Spearman’s correlation analysis confirmed that completion rates for surveys were not associated with any demographic variables (all p values ≥ 0.668) or affective state measures (p values ≥ 0.122).

Consistent with the observation that SSRIs typically act over 4-6 weeks to produce symptom-level change, drug effects on self-report affective measures (Patient Health Questionnaire, Generalized Anxiety Disorder scale, and Apathy Motivation Index) were not detectable during the study (Table 1).^21–23^ This is in line with previous studies in both healthy and clinical participants, providing the basis for testing whether EMA can act as an early and more sensitive readout of affective changes.^11,24^

### Momentary mood reflects distinct contributions of reward outcome and reward prediction error

In support of computational accounts of mood, mixed effect analyses confirmed that positive affect (PA) was dynamically modulated by both reward outcomes and RPEs, with these quantities independently predicting positive affect dynamics. As expected, RPEs and reward outcomes each drove positive affect (rt: β = 0.27, p < 0.001; RPEt: β = 0.04, p = 0.029; Supplementary Table 1), over-and-above prior positive affect (PAt-1: β = 0.33, p < 0.001) and time of activity (Time of Dayt: β = 0.05, p = 0.009).

### SSRIs amplify the impact of reward outcome and RPEs on momentary mood

Mixed-effect analyses confirmed that citalopram led to an increase in overall levels of positive affect over the course of the study (Time in Studyt*SSRI: β = 0.08, p = 0.025; Figure 2C), particularly over the late-study phase from days 9-11 (t(51) = -3.04, p = 0.004; Supplementary Figure 1).

**Figure 2.**
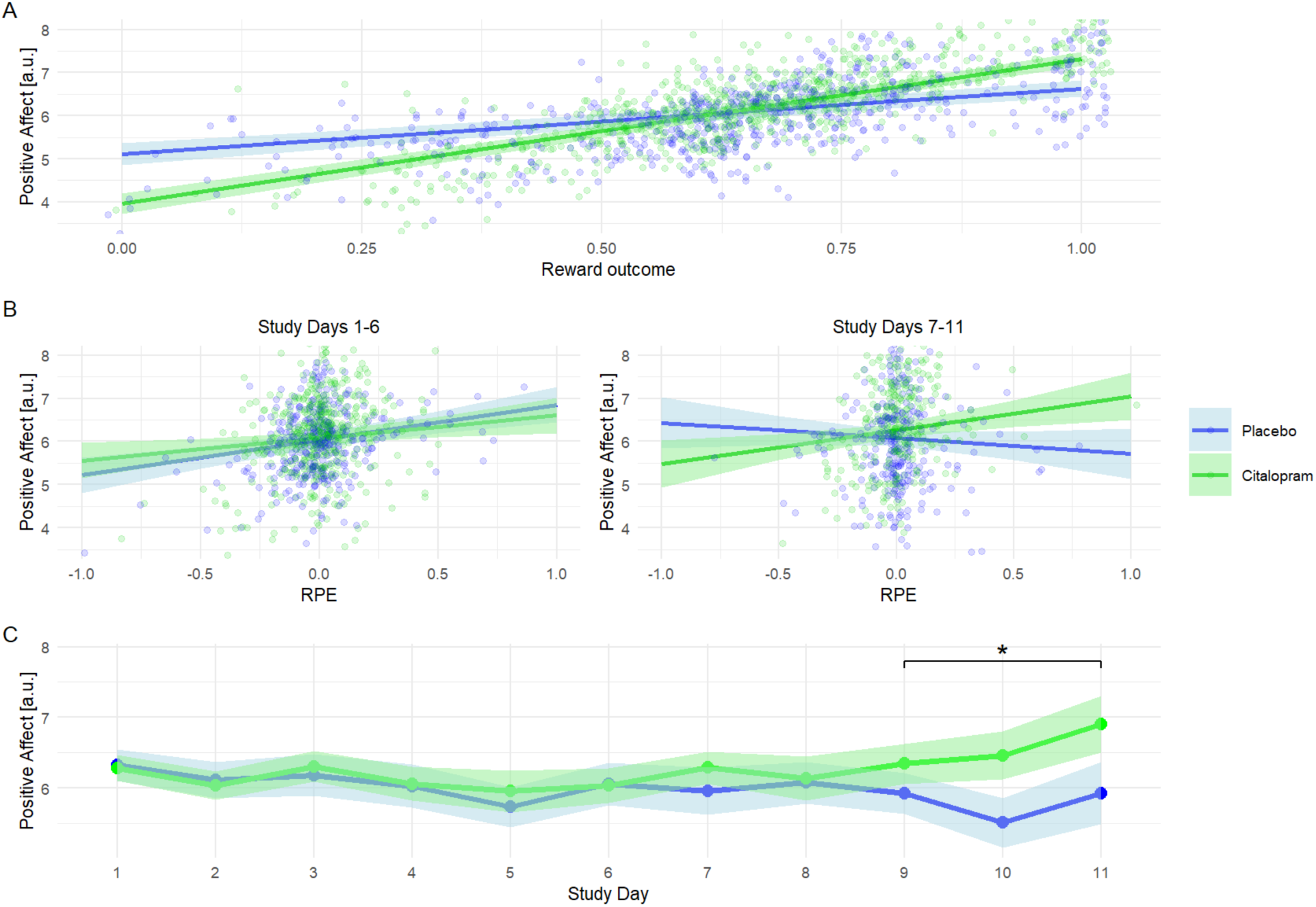
Citalopram amplifies the impact of reward outcome on mood and more gradually amplifies the impact of reward prediction errors over time. Model-predicted PA response to **(A) reward outcome** (i.e. post-activity enjoyment; stronger effect in citalopram group), **(B) reward prediction error** (i.e. degree to which the activity was more enjoyable than expected; stronger increase over time in citalopram group; linear effect split into study days 1-6 and 7-11 for visualisation purposes), and **(C) Time in Study** (greater increase in positive affect in citalopram group, particularly over study days 9-11; linear effect plotted as mean within-participant PA per study day for visualisation purposes). Bands for each line represent the SEM. Raw (non z-scored) data are plotted for interpretability.

Crucially, these improvements in momentary mood occurred in the context of larger changes in the underlying reward processes we proposed (Table 2). Citalopram significantly amplified the impact of reward outcome on positive affect (rt*SSRI: β = 0.20, p < 0.001; Figure 2A). Moreover, citalopram gradually enhanced the impact of RPEs on positive affect, across the study duration (RPEt*SSRI*Time in Studyt: β = 0.13, p < 0.001; Figure 2B). These findings indicate that citalopram’s effects on mood-reward dynamics emerge on a much shorter timescale than the conventional 4 to 6 weeks required for depressive symptom reduction.^25,26^

**Table 2.** Estimated effects of SSRI, Reward Outcome, Reward Prediction Error, and Time in Study on positive affect.

| Parameter | Effect | Standard Error | t | Degrees of freedom | p-value |
| --- | --- | --- | --- | --- | --- |
| Intercept | -0.01 | 0.08 | -0.15 | 55 | 0.882 |
| PA<sub>t-1</sub> | **0.33** | 0.04 | 9.36 | 1230 | **<0.001** |
| r<sub>t</sub> | **0.16** | 0.03 | 5.19 | 1304 | **<0.001** |
| RPE<sub>t</sub> | 0.03 | 0.03 | 1.14 | 1287 | 0.256 |
| SSRI | 0.03 | 0.11 | 0.23 | 55 | 0.821 |
| Time of Day<sub>t</sub> | **0.05** | 0.02 | 3.04 | 1260 | **0.002** |
| Time in Study<sub>t</sub> | -0.05 | 0.03 | -1.78 | 1286 | 0.075 |
| PA<sub>t-1</sub> * SSRI | -0.01 | 0.05 | -0.25 | 1262 | 0.800 |
| r<sub>t</sub> * SSRI | **0.20** | 0.04 | 4.46 | 1306 | **<0.001** |
| RPE<sub>t</sub> * SSRI | 0.02 | 0.04 | 0.52 | 1286 | 0.605 |
| Time in Study<sub>t</sub> * SSRI | **0.08** | 0.04 | 2.24 | 1286 | **0.025** |
| Time in Study<sub>t</sub> * PA<sub>t-1</sub> | 0.03 | 0.03 | 1.11 | 1294 | 0.266 |
| Time in Study<sub>t</sub> * r<sub>t</sub> | -0.02 | 0.03 | -0.71 | 1268 | 0.479 |
| Time in Study<sub>t</sub> * RPE<sub>t</sub> | **-0.07** | 0.03 | -2.57 | 1276 | **0.010** |
| PA<sub>t-1</sub> * SSRI * Time in Study<sub>t</sub> | 0.05 | 0.04 | 1.19 | 1293 | 0.236 |
| r<sub>t</sub> * SSRI * Time in Study<sub>t</sub> | 0.001 | 0.04 | 0.02 | 1275 | 0.986 |
| RPE<sub>t</sub> * SSRI * Time in Study<sub>t</sub> | **0.13** | 0.04 | 3.38 | 1267 | **<0.001** |

**Note:** PA<sub>t-1</sub> = pre-activity positive affect; r<sub>t</sub> = post-activity reward outcome; RPE<sub>t</sub> = post-activity Reward Prediction Error; \* = interaction; **bold** indicates significance at p<0.05.

Follow-up analyses probed the valence specificity of this effect, finding that it was driven by positive RPEs (+RPEt*SSRI*Time in Studyt: β = 0.10, p = 0.005; Supplementary Table 2) and not negative RPEs (p = 0.158), consistent with our predictions based on computational modelling results in laboratory experiments.^11^

To examine the time course of these affect-reward changes across the study, a supplementary analysis was conducted using a two-stage mixed-effects modelling approach. Here, stronger RPE-affect coupling during the dosing period (study days 1-8) predicted higher levels of PA during washout (Participant-specific RPE slope (days 1-8): β = 2.33, p = 0.035; Supplementary Table 3). This preliminary evidence indicates that later increases in PA may be attributed to earlier changes in reward–affect dynamics during SSRI dosing.

### SSRIs decrease the impact of negative mood on reward expectation

Consistent with previous research on affective bias, momentary positive and negative affect (NA) acted as independent and opposing moderators of reward expectation.^8^ As expected, reward expectation was higher as a function of current positive affect and lower as a function of negative affect (PAt: β = 0.22, p < 0.001; NAt: β = -0.23, p < 0.001; Table 3).

**Table 3.** Estimated effects of SSRI, positive and negative affect, lagged Reward Prediction Errors, and Time in Study on reward expectation.

| Parameter | Effect | Standard Error | t | Degrees of freedom | p-value |
| --- | --- | --- | --- | --- | --- |
| Intercept | 0.04 | 0.08 | 0.42 | 61 | 0.677 |
| PA<sub>t</sub> | **0.22** | 0.05 | 4.00 | 621 | **<0.001** |
| NA<sub>t</sub> | **-0.23** | 0.06 | -3.92 | 607 | **<0.001** |
| RPE<sub>t-1</sub> | -0.07 | 0.04 | -1.76 | 1090 | 0.079 |
| SSRI | -0.05 | 0.11 | -0.45 | 61 | 0.657 |
| Time of Day<sub>t</sub> | -0.15 | **0.03** | -5.70 | 1081 | **<0.001** |
| Time in Study<sub>t</sub> | **-0.10** | 0.04 | -2.39 | 1103 | **0.017** |
| PA<sub>t</sub> * SSRI | 0.02 | 0.07 | 0.32 | 695 | 0.748 |
| NA<sub>t</sub> * SSRI | **0.21** | 0.08 | 2.73 | 586 | **0.007** |
| RPE<sub>t-1</sub> * SSRI | **0.12** | 0.05 | 2.22 | 1102 | **0.026** |
| Time in Study<sub>t</sub> * SSRI | 0.02 | 0.06 | 0.32 | 1103 | 0.750 |
| Time in Study<sub>t</sub> * PA<sub>t</sub> | -0.004 | 0.04 | -0.09 | 1103 | 0.930 |
| Time in Study<sub>t</sub> * NA<sub>t</sub> | -0.06 | 0.04 | -1.38 | 1094 | 0.169 |
| Time in Study<sub>t</sub> * RPE<sub>t-1</sub> | **-0.08** | 0.04 | -1.99 | 1090 | **0.046** |
| PA<sub>t</sub> * SSRI * Time in Study<sub>t</sub> | -0.03 | 0.06 | -0.43 | 1108 | 0.666 |
| NA<sub>t</sub> * SSRI * Time in Study<sub>t</sub> | 0.02 | 0.06 | 0.31 | 1100 | 0.756 |
| RPE<sub>t-1</sub> * SSRI * Time in Study<sub>t</sub> | **0.11** | 0.05 | 2.10 | 1088 | **0.036** |

**Note:** PA<sub>t</sub> = pre-activity positive affect; NA<sub>t</sub> = pre-activity negative affect; RPE<sub>t-1</sub> = lagged post-activity Reward Prediction Error; \* = interaction; **bold** indicates significance at $p < 0.05$ .

Citalopram selectively attenuated the impact of negative affect on reward expectation (NAt*SSRI: β = 0.21, p = 0.007; Figure 3A), without modulating the increase in reward expectation by positive affect (PAt*SSRI: β = 0.02, p = 0.748; Figure 3B). Contributing to this effect was a significantly higher proportion of floor-level negative affect reports under citalopram relative to placebo (45.9% vs. 37.3%; OR = 0.70, p < 0.001), suggesting that citalopram partially reduced the occurrence of elevated negative affect states in addition to attenuating their impact on reward expectation when they did occur. When these observations were excluded, the interaction between negative affect and drug group was attenuated (NAt*SSRI: β = 0.19, p = 0.080), likely reflecting reduced power following removal of 842 observations (42% of the sample). Together, these findings align with an early correction of negative affective bias under citalopram, as reported in laboratory-based studies, as well as reductions in overall levels of negative affect.^18,27^

**Figure 3.**
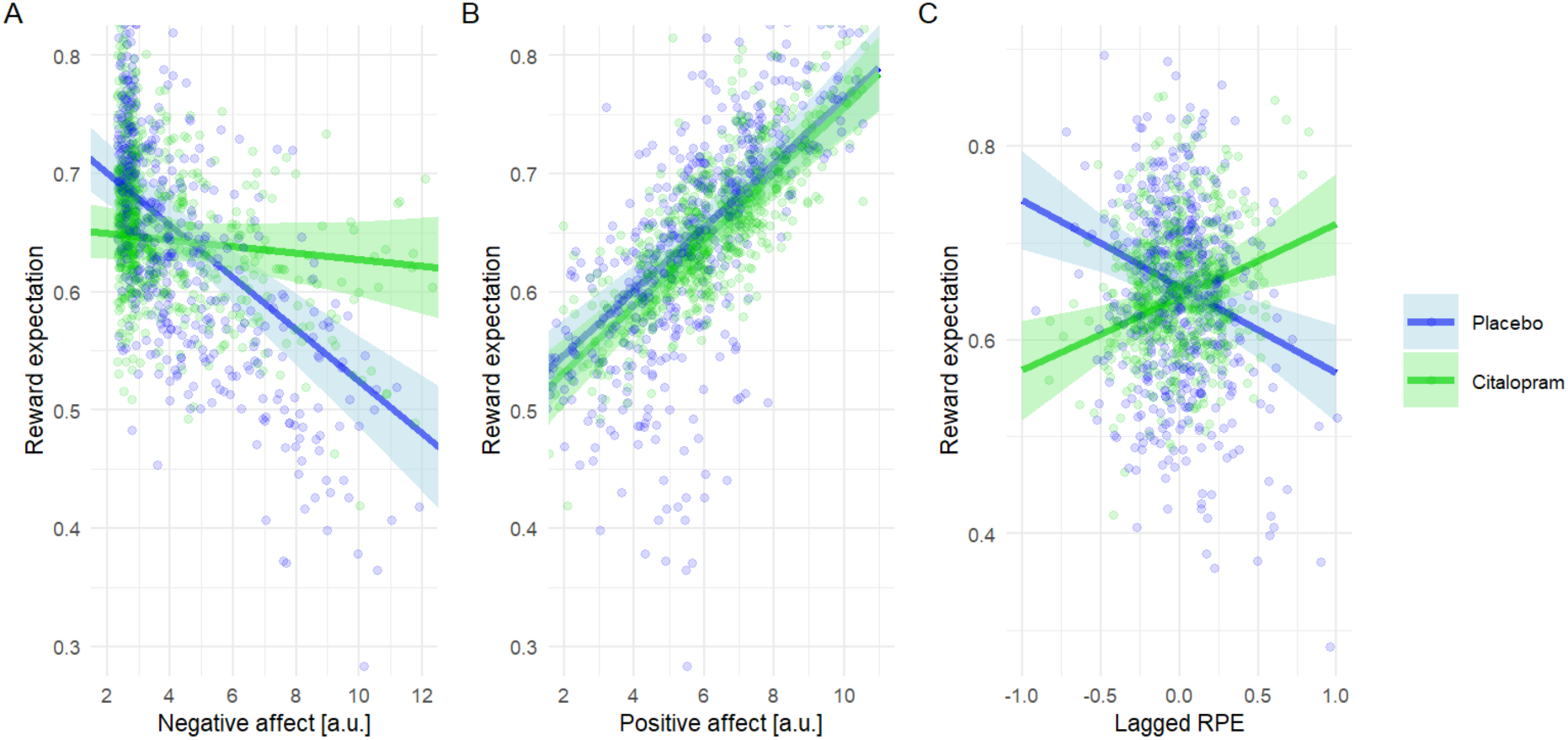
Citalopram attenuates the impact of negative affect on reward expectation while boosting the effect of prior reward prediction errors. Model-predicted reward expectation to **(A) negative affect** (weakened effect in the citalopram group) and **(B) positive affect** (no difference between groups). **(C) Lagged reward prediction errors,** which represent the carryover impact of RPEs from the preceding activity (increased effect in citalopram group but diminished effect in placebo group). Bands for each line represent the SEM. Raw (non z-scored) data are plotted for interpretability.

### SSRIs increase the coupling between prior RPEs and reward expectation

In addition to changes in momentary mood, lagged RPE was also a significant predictor of subsequent reward expectation. Specifically, citalopram boosted the carryover impact of prior activity RPE on current reward expectation (RPEt-1*SSRI: β = 0.12, p = 0.026), such that a better-than-expected prior activity increased reward expectation for the subsequent activity (Figure 3C). By contrast, this carry-over effect was absent - and nominally reversed - in the placebo group, suggesting that citalopram selectively enhanced the degree to which positive prior outcomes shaped prospective reward anticipation. Consistent with the gradual time course of SSRI action, this between-group difference strengthened over the course of the study period (RPEt-1*SSRI*Time in Studyt: β = 0.11, p = 0.036).

Follow-up analyses on SSRI valence specificity suggested that the SSRI-led amplification of lagged RPE on subsequent reward expectation was stronger for positive RPEs (+RPEt-1*SSRI*Time in Studyt: β = 0.10, p = 0.072; Supplementary Table 4) than negative RPEs (p = 0.492). This is directionally consistent with the above finding that SSRIs selectively amplified the impact of positive but not negative RPEs on positive affect (Supplementary Table 2).

## Discussion

Serotonergic antidepressants lead to improvements in mood symptoms that are typically only detectable after weeks or months, despite exerting rapid neurochemical effects on the circuitry underlying affective processing.^2,28^ At the other extreme, computational work suggests that SSRIs may modulate the bidirectional relationship between momentary mood and reward perception, but using monetary tasks that operate on the order of seconds to minutes, leaving it unknown whether these mechanisms translate to real-world contexts and timescales.^11^

In this double-blind, placebo-controlled study, we combined eight days of SSRI administration with eleven days of EMA, to characterize how affect–reward dynamics unfold across daily life. We demonstrate that SSRIs detectably uplift mood by the end of this brief period (Figure 2C) and further find support for affect-reward mechanisms being involved in this improvement. Specifically, we find that SSRIs rapidly upregulate the impact of reward outcome and RPEs on positive affect, while in parallel attenuating the coupling between negative affect and reward expectations. Our findings thus bring together two lines of research into antidepressant drug action. First, in agreement with laboratory-based studies, we show that SSRI effects are more specific than a direct action on mood or reward processes, instead increasing how ‘well-tuned’ mood is to unexpected outcomes (Figure 2B).^11^ Second, our results build on evidence that SSRIs acutely reduce negative affective bias, which studies have previously demonstrated with negative emotional cues (e.g., aversive facial expressions).^7,29^ Our results show that this attenuation in negative bias also operates on real-world reward, shaping how mood informs reward expectations for upcoming activities (Figure 3A). Interestingly, the attenuation in negative bias appears partly driven by lower levels of negative affect under SSRI. Coupled with the gradual upward drift in positive affect, this suggests that the indirect effect of SSRIs on reward–mood coupling may consolidate into a broader shift in affective state. Overall, these complementary mechanisms suggest that SSRIs may promote a more positively weighted feedback loop that operates over daily life, potentially ameliorating deficits in these processes in depression, thus contributing to the gradual mood improvements typically observed with antidepressant treatment. ^17^

While reward outcome and RPEs are conceptualized as contemporaneous drivers of momentary mood, thought to be predominantly reliant upon dopaminergic signals, our results showed a temporal divergence in their serotonergic modulation.^30^ The SSRI-amplified impact of reward outcome on mood was evident throughout our study, which chimes with human and animal research demonstrating the role of serotonin in reward valuation and affective processing.^31,32^

By contrast, the SSRI-modulation of RPE effects on mood developed progressively with continued dosing. This latter pattern may reflect more complex interactions between serotonergic and dopaminergic systems, and points to a gradual retuning of affect-reward associations over time.^33^ We speculate that this mechanism may drive the later increases we observed in overall mood, as supported by our supplementary analysis linking the strength of pre-washout RPE-affect coupling to post-washout PA. This possibility could be tested by future work, potentially using continuous-time models applied to EMA studies over a longer duration of time.^34^ If confirmed, this could provide a valuable assay for more rapidly evaluating the effects of SSRIs.

In keeping with computational frameworks, our results support the influential momentum-based model of mood.^8^ Specifically, when activities were better than expected, SSRIs promoted the carryover of this signal to reward expectations about subsequent activities (Figure 3C). While previous EMA studies have demonstrated that exam grade prediction errors update expectations about subsequent exams, our design allowed us to demonstrate such updating generalizes across highly distinct goal-directed activities (e.g., going for a walk, eating a meal).^35,36^ We interpret this carryover as an emergent property of ‘mood as momentum’: affective information generalizes across contexts and persists over time. In this way, a positive experience on a walk increases expectations for an entirely different activity, like eating a meal, despite no direct relevance between them. This mechanism is strikingly consistent with the rationale underlying behavioral activation in cognitive behavioral therapy (CBT) for depression, which encourages patients to enhance both the quantity and quality of the rewarding, goal-directed activities they engage with throughout the day.^37^ An unexpected finding was that in the placebo group, the carryover effect instead became progressively negative over time. This could reflect an EMA “training” effect, whereby repeated reflection on mood and expectations encourages the use of a regression-to-the-mean heuristic, (e.g., ‘the previous activity went much better than expected, so I expect the next activity to not be as good’).^38,39^ Given that this drift would be expected to produce more pessimistic expectations, its attenuation by SSRIs is intriguing, and may explain why the SSRI group evidenced gains in overall mood compared to placebo by the end of the study (Figure 2C).

Our findings link the mechanisms of antidepressant action to valence-dependent RPE signaling in depression. Specifically, an EMA study reported that individuals with heightened depressive symptoms displayed blunted affective responses to positive RPEs, while responses to negative RPEs were comparable to those without depression.^36^ Our results mirror this pattern, with SSRI-induced changes in mood-reward dynamics emerging specifically for positive, but not negative, RPEs. This valence-specificity, dampened in depression and selectively amplified by SSRIs, suggests a shared computational substrate linking serotonergic modulation and affective dysfunction. Although our findings are limited by the non-clinical nature of our participants, they nonetheless highlight positive RPE processing as a potential target for behavioral interventions, which may promote the reframing and reinforcement of positive experiences.

## Conclusions

In conclusion, our study provides a novel lens into the early effects of SSRIs on affective dynamics in daily life. We show that serotonin modulates both the immediate and carryover effects of consummatory reward signals onto mood and subsequent expectations, thereby offering a more nuanced understanding of the complex interplay between these processes. Moreover, we provide new insight into the mechanistic targets of antidepressant treatment and their potential early detection using real-world data, helping to bridge the gap between the basic neurobiological mechanisms of SSRIs and clinical symptom change in depression.

## Methods

The hypotheses and analysis plan for this study were pre-registered on Open Science Framework (https://osf.io/92rq6/overview).^40^ Deviations from the pre-registration are reported in Supplementary Table 5.^41^

### Participants

70 healthy participants, matched by age and gender across drug groups, were recruited for this double-blind, placebo-controlled study. Eligibility was assessed in two stages, initially by an online questionnaire battery and then by a medical doctor, who completed a structured medical history during the in-person study session on Day 1. Inclusion criteria were: 18 to 40 years of age, no current health issues, no history of neurological or psychiatric disorders, no prior allergic reaction to any drug, no regular medication usage, and an electrocardiogram confirming the absence of QT interval prolongation. The study protocol was approved by the University College London research ethics committee (Ethics ID: 19601/001), with informed consent obtained from all participants.

### Pharmacological procedure

40 participants were randomly allocated to receive a daily oral dose of the SSRI citalopram (20 mg), while the remaining 30 participants were allocated to placebo. Participants were informed of the double-blind design of the study and were instructed to take their medication for eight consecutive days.

The initial dose was taken at the in-person study session on Day 1, following completion of mood and psychiatric symptom questionnaires and baseline testing. For subsequent days, participants were instructed to take their medication at home at a similar time of day, typically with breakfast. Participants attended a follow-up study session on Day 8, the last day of drug administration. A three-day washout period followed, spanning Day 9 to Day 11, to probe sustained effects beyond the dosing phase.

During study debrief, subjective drug effects and group allocation guesses were captured. Of the participants that received citalopram, 28 (70%) experienced minor side effects, while 27 (67%) correctly guessed that they were given SSRI. 19 participants (63%) correctly guessed that they had received placebo (Supplementary Table 6). Group allocation did not affect drop-out at the in-person study session on Day 8 (odds ratio = 1.0, p = 1.00).

### Instruments

To assess potential drug effects on affective state measures, the Patient Health Questionnaire (PHQ-9), Generalized Anxiety Disorder scale (GAD-7), and Apathy Motivation Index (AMI) were administered to participants during the in-person study sessions on Days 1 and 8.^21–23^ The PHQ-9 and GAD-7 were chosen as established self-report measures of depressive and anxiety symptoms, respectively, whereas the AMI was selected as a measure of sub-clinical levels of apathy. Additional questionnaires on psychiatric symptoms and personality traits were administered to participants during the in-person study sessions (not analyzed here).

### EMA procedure

Participants completed eleven days of EMA using the ‘The Happiness Project’ smartphone app (https://rutledgelab.org/), which was installed on participants’ phones during the study session on Day 1. They were expected to complete four pre-activity surveys and four post-activity surveys per day, incentivized by a bonus payment proportional to their EMA survey completion rate, on top of a base rate of pay. Daily notifications for the pre-activity survey were scheduled at 9:00am, 12:00pm, 3:00pm and 6:00pm, and surveyed momentary mood and a rewarding event assessment. Importantly, each of these surveys generated a paired post-activity survey, yielding up to eight daily probes of momentary mood and reward.

Momentary mood was measured using an adapted version of Mood Zoom, a 6-item measure developed to assess dynamic mood states in clinical and healthy populations.^42^ Participants were prompted to rate how “Anxious”, “Sad”, “Angry”, “Irritable”, “Energetic”, and “Happy” they were currently feeling on a 7 point-Likert scale, ranging from 1 (“Not at all”) to 7 (“Very much”).

For reward event assessment, participants were asked to identify a salient activity that would occur in the next two hours (“What are you most excited or most nervous about doing?”). They received extensive instructions from the researcher about identifying ‘SMART’ activities, defined as specific, measurable, achievable, relevant, and time bound. Participants were also asked to select activities where they were unsure of the outcome, and where possible to avoid re-using the same types of activities. Representative examples of participant activities included “walking the dog”, “getting lunch with a friend” and “giving a presentation”. After identifying an activity, participants were asked to assign it to one of the following categories: “working/studying”, “household chores”, “eating/drinking”, “social activities”, “active leisure”, “passive leisure”, “resting/napping”, “browsing computer/phone”, “daydreaming/mind-wandering”, and “other”. Recognizing that some activities fall into multiple categories, participants were instructed to select the category that best reflected the most salient aspect of the activity. To trigger the paired post-activity survey, participants were then asked to identify the time that the activity would be finished. The app dynamically programmed a notification based on this time. Finally, participants rated the activity’s predicted reward across four reward-related dimensions, using a slider from 0-100 with anchors “Not at all” and “Very much” (i.e., “How *enjoyable* do you think the activity will be?”, “How much *sense of achievement* will the activity bring you?”, “How *connected to others* will the activity make you feel?”, and “How *effortful* will the activity be?”).

The post-activity survey was notified at the time specified by the participant, and assessed momentary mood as above, followed by probes for the actual experience of reward, measured along the same reward-related dimensions and sliding scale (i.e., “How *enjoyable* did you find the activity?”, “How much *sense of achievement* did the activity bring you?”, “How *connected to others* did the activity make you feel?”, and “How *effortful* was the activity?”; 0-100).

### EMA response processing

Participants who dropped out of the study after the in-person session on Day 1 were removed from the dataset, leaving a final analysis sample of 66 participants. As recommended by Jaso et al., surveys in the final analysis sample were checked for careless responding (i.e., <1S or >60s time spent on each survey item).^43^ This led to the removal of 7.55% of paired surveys. The average survey duration following EMA cleaning was 37.3s (SD = 16.1).

### Momentary mood

The latent variable structure of momentary mood was determined using parallel and exploratory factor analysis.^44^ In line with prior studies, this yielded a two-factor solution representing positive affect (PA) and negative affect (NA).^42^ As expected, the items “Happy” and “Energetic” loaded preferentially onto the PA factor, while “Anxious”, “Sad”, “Angry”, and “Irritable” loaded preferentially onto the NA factor. Items with weights less than 0.4 were not applied when calculating each factor (Supplementary Table 7).^45^

Both PA and NA factors showed acceptable internal consistency given the inherently dynamic nature of momentary affect (Cronbach’s α = 0.74 for NA, 0.66 for PA). Supporting construct validity, PA was confirmed to have substantial temporal stability across the study (r = 0.90 between first and second-half study means; ICC = 0.47), as did NA (r = 0.91; ICC = 0.58).

### Reward

Of the four reward items that were captured by EMA, ‘enjoyment’ was selected as the focus of the current paper because of its relevance across participant activities.

To evaluate reward across multiple dimensions, parallel and exploratory factor analysis was also performed on all reward items. This yielded a two-factor solution representing pleasure and mastery, with ‘enjoyment’ and ‘social connectedness’ loading onto pleasure, and ‘achievement’ and ‘effort’ loading onto mastery. Full analysis plans and results for pleasure and mastery are reported in the Supplementary Information (Supplementary Table 8 and 9).

### Reward Prediction Errors

RPEs were calculated as the difference between the actual experience of enjoyment (reward outcome, r) and the expected enjoyment (reward prediction, ν) extracted from paired post- and pre-activity surveys, respectively.

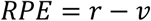

### Planned analyses

Given the hierarchical structure of the dataset (i.e., mood and reward ratings nested within participants and within groups), multilevel regression models were used to account for participant-specific and SSRI group-specific effects. Linear mixed models were specified and tested using the “lme4” package in RStudio (R v.4.4.0).^46^ To account for linear time effects during the study, the number of days elapsed since the first survey was calculated, ranging from study days 1 to 11. To account for non-linear time effects, due to the diurnal modulation of positive affect in relation to the sleep-wake cycle, a regressor was calculated that quantified the number of hours that the current survey fell from the midpoint of the sleep-wake cycle as an inverted U-shaped quadratic.^47,48^ All models therefore included the variables ‘Time in Study’ and ‘Time of Day’, as well as random intercepts for participants.

Continuous variables were assessed for normality. Right-skewed variables (pre- and post-survey negative affect) were log-transformed, while a cube-root transformation was applied to Time of Day to correct for left skewness. All continuous variables were subsequently converted to z-scores before analyses, in line with best practices and to obtain standardized beta coefficients.^49^

Finally, to assess multicollinearity, average within-participant correlations for RPE, r and ν were calculated, in addition to variance inflation factors (VIFs) for fixed effects in all models. As expected, RPE and r were moderately positively correlated (r = 0.39), whereas RPE and ν were moderately negatively correlated (r = -0.48). All VIF values were below 2.99, suggesting acceptable levels of collinearity.

### Impact of reward outcomes and RPEs on momentary mood

To examine RPEs, reward outcomes, and pre-activity positive affect as drivers of momentary mood, post-activity positive affect was modelled as a function of RPEt, rt and PAt-1. Three-way interactions with drug group (placebo, citalopram) and Time in Study captured potential SSRI effects evolving over time:

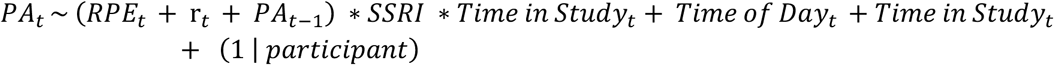

where t represents EMA survey timepoint. To test whether associations with SSRI usage are dependent on RPE valence, this model was rerun with positive and negative RPEs entered as separate terms.

### Coupling of reward expectations with momentary mood

Pre-activity reward expectation was regressed onto pre-activity positive and negative affect. Lagged RPEs (derived from the previous activity at timepoint t-1) were included in the model as a representation of reward momentum, to determine whether RPEs from prior activities exert a carryover effect onto future reward expectation.^8^ As before, interactions with drug group and Time in Study captured potential SSRI effects on reward expectations over time:

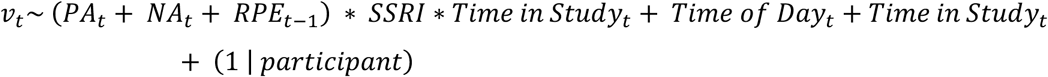

As a follow-up analysis, this model was also rerun with positive and negative RPEs entered as separate terms.

## Data availability

The data that support the findings of this study are available from the corresponding author upon reasonable request.

## Acknowledgements

This work was supported by a Medical Research Council Clinician Scientist Fellowship awarded to L.M. (MR/S006613/1), as well as a Medical Research Council Career Development Award (MR/N02401X/1), a NARSAD Young Investigator Grant (27674) from the Brain and Behavior Research Foundation, P and S Fund, and a National Institute of Mental Health grant (1R01MH124110) awarded to R.B.R..

We would also like to acknowledge Tom Metherell (University College London) for his conceptual input and contributions to the development of analyses and pre-registration document during his PhD rotation.

## Author contributions

R.M. analysed the data and wrote the manuscript. M.C. and A.D. collected the data. L.M. conceived and designed the study, advised on analysis, and wrote the manuscript. R.B.R. conceived and designed the study, advised on analysis, and contributed to manuscript preparation. All authors reviewed and approved the final manuscript.

## Competing interests

The authors declare no competing interests.

## Notes

### Competing Interest Statement

The authors have declared no competing interest.

