## Supplementary_Material for "Serotonin reuptake inhibition rapidly enhances affect-reward coupling and improves mood over time"

[†] Current affiliation: Department of Psychology, Bath University, Bath, UK

#### Supplementary Information

**Supplementary Table 1. Estimated effects of prior positive affect, Reward Prediction Errors and Reward Outcome on positive affect**

| $PA_t \sim RPE_t + r_t + PA_{t-1} + Time\ of\ Day_t + Time\ in\ Study_t + (1 \mid participant)$ | | | | | |
|---|---|---|---|---|---|
| **Parameter** | **Effect** | **Standard Error** | ***t*** | **Degrees of freedom** | **p-value** |
| Intercept | -0.001 | 0.06 | -0.01 | 57 | 0.993 |
| $PA_{t-1}$ | **0.33** | 0.02 | 13.70 | 1280 | **<0.001** |
| $r_t$ | **0.27** | 0.02 | 11.86 | 1318 | **<0.001** |
| $RPE_t$ | **0.04** | 0.02 | 2.619 | 1297 | **0.029** |
| Time of $Day_t$ | **0.05** | 0.02 | 2.62 | 1270 | **0.009** |
| Time in $Study_t$ | 0.005 | 0.02 | 0.31 | 1196 | 0.760 |

***Note:*** *$PA_{t-1}$ = pre-activity positive affect; $r_t$ = post-activity reward outcome; $RPE_t$ = post-activity Reward Prediction Error; * = interaction;* ***bold*** *indicates significance at p<0.05.*

### Supplementary Information

#### SSRIs improve momentary mood from early to late study phases

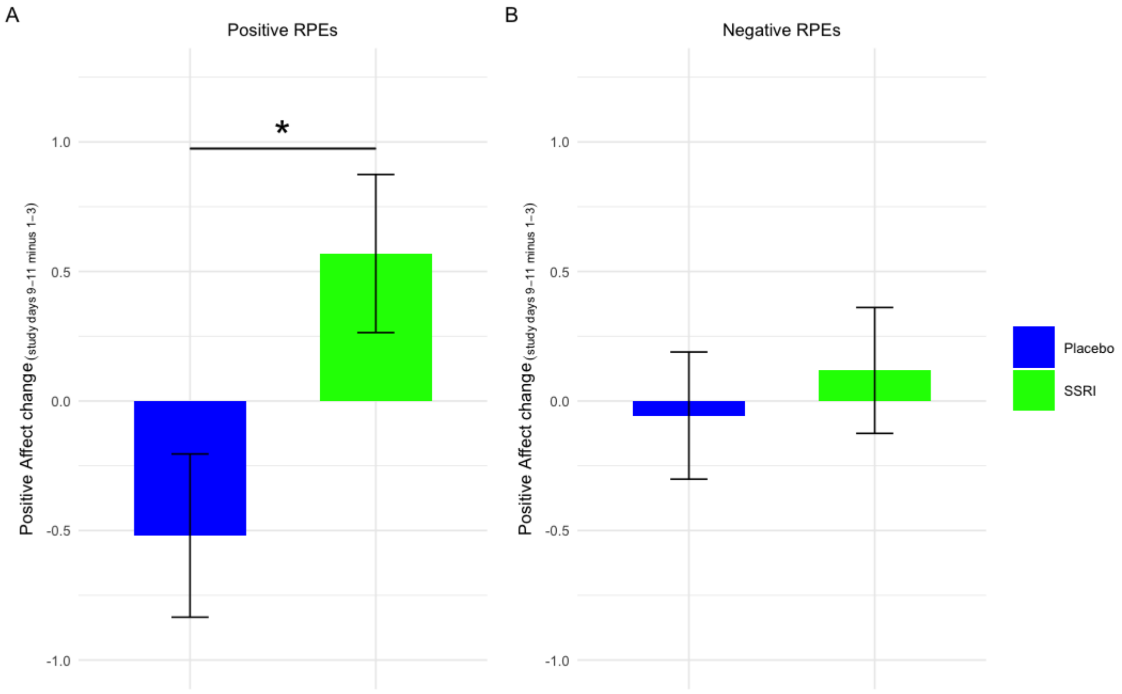

**Supplementary Figure 1. Citalopram increases positive affect from early to late study phases.** Change in mean (z-scored) positive affect between study days 9-11 and 1-3, split for (A) positive RPEs (significant increase in citalopram group) and (B) negative RPEs (no difference between groups). Error bars indicate SEM. * indicates significance at $p<0.05$. Raw (non z-scored) data are plotted for interpretability.

Citalopram significantly increased overall levels of positive affect between early (days 1–3) and late (days 9–11) study phases ($t_{(51)} = -3.04$, $p = 0.004$). Follow-up analyses found that this effect was linked specifically to changes in mood following positive real-world RPEs ($t_{(41)} = -2.48$, $p = 0.017$; Supplementary Figure 1A) and not changes in mood following negative RPEs ($t_{(43)} = -0.50$, $p = 0.617$; Supplementary Figure 1B), consistent with our predictions based on computational modelling results in laboratory experiments.[1]

#### Supplementary Information

**Supplementary Table 2. Estimated effects of SSRI, prior positive affect, positive Reward Prediction Errors, negative Reward Prediction Errors, and Reward Outcome on positive affect**

| $PA_t \sim (+RPE_t + -RPE_t + r_t + PA_{t-1}) * Group * Time\ in\ Study + Time\ of\ Day_t + Time\ in\ Study_t + (1 \mid participant)$ | | | | | |
|---|---|---|---|---|---|
| **Parameter** | **Effect** | **Standard Error** | ***t*** | **Degrees of freedom** | **p-value** |
| Intercept | -0.01 | 0.08 | -0.15 | 55 | 0.878 |
| $PA_{t-1}$ | **0.33** | 0.04 | 9.37 | 1227 | **<0.001** |
| $r_t$ | **0.16** | 0.03 | 4.88 | 1299 | **<0.001** |
| $+RPE_t$ | -0.01 | 0.03 | 0.39 | 1291 | 0.696 |
| $-RPE_t$ | 0.04 | 0.03 | 1.16 | 1273 | 0.248 |
| SSRI | 0.03 | 0.11 | 0.24 | 55 | 0.814 |
| Time of $Day_t$ | **0.05** | 0.02 | 2.92 | 1257 | **0.004** |
| Time in $Study_t$ | -0.05 | 0.03 | -1.83 | 1283 | 0.067 |
| $PA_{t-1}$ * SSRI | -0.01 | 0.05 | -0.27 | 1260 | 0.789 |
| $r_t$ * SSRI | **0.20** | 0.05 | 4.36 | 1303 | **<0.001** |
| $+RPE_t$ * SSRI | 0.001 | 0.04 | 0.02 | 1289 | 0.983 |
| $-RPE_t$ * SSRI | 0.02 | 0.04 | 0.39 | 1298 | 0.700 |
| Time in $Study_t$ * SSRI | **0.08** | 0.04 | 2.28 | 1283 | **0.023** |
| Time in $Study_t$ * $PA_{t-1}$ | 0.03 | 0.03 | 1.11 | 1290 | 0.265 |
| Time in $Study_t$ * $r_t$ | -0.05 | 0.03 | -0.78 | 1264 | 0.434 |
| Time in $Study_t$ * $+RPE_t$ | **-0.05** | 0.03 | -1.99 | 1255 | **0.046** |
| Time in $Study_t$ * $-RPE_t$ | -0.04 | 0.03 | -1.22 | 1272 | 0.221 |
| $PA_{t-1}$ * SSRI * Time in $Study_t$ | 0.05 | 0.04 | 1.20 | 1290 | 0.230 |
| $r_t$ * SSRI * Time in $Study_t$ | 0.006 | 0.04 | 0.14 | 1271 | 0.888 |
| $+RPE_t$ * SSRI * Time in $Study_t$ | **0.10** | 0.04 | 2.84 | 1266 | **0.004** |
| $-RPE_t$ * SSRI * Time in $Study_t$ | 0.06 | 0.04 | 1.41 | 1265 | 0.158 |

***Note:*** *$PA_{t-1}$ = pre-activity positive affect; $r_t$ = post-activity reward outcome; $+RPE_t$ = post-activity positive Reward Prediction Error; $-RPE_t$ = post-activity negative Reward Prediction Error; * = interaction;* ***bold*** *indicates significance at $p<0.05$.*

### Supplementary Information

##### Estimation of RPE-affect coupling during and after SSRI dosing

To examine the time course of affect-reward coupling during and after SSRI dosing, a two-stage mixed-effects modelling approach was used. In the dosing period (i.e., study days 1-8), we estimated participant-specific random slopes, capturing the interaction between RPE and Time in Study in a model predicting dosing-PA. This allowed us to index individual differences in RPE-affect coupling during SSRI administration.

$$\boldsymbol{Dosing\ PA_t \sim Dosing\ PA_{t-1} + Time\ of\ Day_t + (1 + RPE_t * Time\ in\ Study_t \mid participant)}$$

These slopes, along with mean dosing-PA, were then used as participant-level predictors of momentary positive affect during the washout period (i.e., study days 9-11). This approach enabled us to test whether earlier reward–affect dynamics prospectively predicted later positive affect.

$$\boldsymbol{Washout\ PA_t \sim \left(Mean\ dosing - PA_{(days\ 1-8)} * \ Participant - specific\ RPE\ slope_{(days\ 1-8)}\right) + Time\ of\ Day_t + (1 \mid participant)}$$

**Supplementary Table 3. Estimated effects of mean dosing positive affect and RPE-affect coupling on washout positive affect**

| $\boldsymbol{Washout\ PA_t \sim \left(Mean\ dosing - PA_{(days\ 1-8)} * \ Participant - specific\ RPE\ slope_{(days\ 1-8)}\right) + Time\ of\ Day_t + (1 \mid participant)}$ | | | | | |
|---|---|---|---|---|---|
| **Parameter** | **Effect** | **Standard Error** | ***t*** | **Degrees of freedom** | **p-value** |
| Intercept | 0.08 | 0.07 | 1.17 | 39 | 0.251 |
| Mean dosing-PA$_{(days\ 1\text{-}8)}$ | **1.05** | 0.08 | 13.1 | 33 | **<0.001** |
| Participant-specific RPE slope$_{(days\ 1\text{-}8)}$ | **2.33** | 1.06 | 2.19 | 34 | **0.035** |
| Time of Day$_t$ | -0.01 | 0.04 | -0.32 | 268 | 0.750 |
| Mean dosing-PA$_{(days\ 1\text{-}8)}$ * Participant-specific RPE slope$_{(days\ 1\text{-}8)}$ | 2.20 | 1.57 | 1.40 | 33 | 0.171 |

***Note:*** *Mean dosing-PA$_{(days\ 1\text{-}8)}$ = average post-activity positive affect during dosing; Participant-specific RPE slope$_{(days\ 1\text{-}8)}$ = individual estimates of RPE-affect coupling during dosing; * = interaction;* ***bold*** *indicates significance at p<0.05.*

Washout positive affect was strongly predicted by mean dosing positive affect (Mean dosing-PA $_{(days\ 1\text{-}8)}$: β = 1.05, p < 0.001), indicating substantial stability in affective levels during and after SSRI administration. Notably, individual estimates of RPE–affect coupling during dosing also significantly predicted washout positive affect (Participant-specific RPE slope $_{(days\ 1\text{-}8)}$: β = 2.33, p = 0.035). Here, individuals with stronger RPE-linked affective responses during the dosing period showed higher washout PA. This preliminary evidence indicates that later increases in positive affect may be partially attributed to earlier changes in reward–affect dynamics during the dosing phase.

### Supplementary Information

**Supplementary Table 4. Estimated effects of SSRI, lagged positive Reward Prediction Errors, lagged negative Reward Prediction Errors, and positive and negative affect on reward expectation**

| $v_t \sim (+RPE_{t-1} + -RPE_{t-1} + PA_t + NA_t) * Group * Time\ in\ Study_t + Time\ of\ Day_t + Time\ in\ Study_t + (1 \mid participant)$ | | | | | |
|---|---|---|---|---|---|
| **Parameter** | **Effect** | **Standard Error** | ***t*** | **Degrees of freedom** | **p-value** |
| Intercept | 0.03 | 0.08 | 0.41 | 60 | 0.683 |
| $PA_t$ | **0.21** | 0.05 | 3.95 | 613 | **<0.001** |
| $NA_t$ | **-0.23** | 0.06 | -3.92 | 607 | **<0.001** |
| $+RPE_{t-1}$ | -0.05 | 0.04 | -1.36 | 1104 | 0.175 |
| $-RPE_{t-1}$ | -0.02 | 0.04 | -0.62 | 1083 | 0.539 |
| SSRI | -0.05 | 0.11 | -0.44 | 60 | 0.661 |
| Time of $Day_t$ | **-0.15** | 0.03 | -5.71 | 1077 | **<0.001** |
| Time in $Study_t$ | **-0.10** | 0.04 | -2.40 | 1099 | **0.016** |
| $PA_t$ * SSRI | 0.03 | 0.07 | 0.35 | 695 | 0.726 |
| $NA_t$ * SSRI | **0.22** | 0.08 | 2.72 | 597 | **0.007** |
| $+RPE_{t-1}$ * SSRI | 0.09 | 0.06 | 1.48 | 1103 | 0.139 |
| $-RPE_{t-1}$ * SSRI | 0.06 | 0.06 | 1.01 | 1103 | 0.315 |
| Time in $Study_t$ * SSRI | 0.02 | 0.06 | 0.33 | 1099 | 0.743 |
| Time in $Study_t$ * $PA_t$ | -0.007 | 0.04 | -0.17 | 1097 | 0.863 |
| Time in $Study_t$ * $NA_t$ | -0.06 | 0.04 | -1.36 | 597 | 0.175 |
| Time in $Study_t$ * $+RPE_{t-1}$ | **-0.08** | 0.04 | -1.99 | 1075 | **0.046** |
| Time in $Study_t$ * $-RPE_{t-1}$ | -0.02 | 0.04 | -0.40 | 1086 | 0.681 |
| $PA_t$ * SSRI * Time in $Study_t$ | -0.02 | 0.06 | -0.37 | 1104 | 0.708 |
| $NA_t$ * SSRI * Time in $Study_t$ | 0.02 | 0.06 | 0.31 | 1096 | 0.761 |
| $+RPE_{t-1}$ * SSRI * Time in $Study_t$ | 0.100 | 0.06 | 1.80 | 1089 | 0.072 |
| $-RPE_{t-1}$ * SSRI * Time in $Study_t$ | 0.04 | 0.06 | 0.69 | 1086 | 0.482 |

***Note:*** *$PA_t$ = pre-activity positive affect; $NA_t$ = pre-activity negative affect; $+RPE_{t-1}$ = lagged post-activity positive Reward Prediction Error; $-RPE_{t-1}$ = lagged post-activity negative Reward Prediction Error; * = interaction;* ***bold*** *indicates significance at $p<0.05$.*

#### Supplementary Information

**Supplementary Table 5. Pre-registration deviations.** Hypotheses 1a and 1b, involving computational modelling, were not analysed due to time and resource constraints. Deviations from the pre-registered analysis plan for Hypotheses 2a, 2b, and 2c are described below.

<table>
<tr><td colspan="4">Deviations to all models</td></tr>
<tr><td colspan="4">Attempts to fit models with random slopes for affect, outcome, and RPE resulted in convergence warnings and singular fits; random intercept-only structures were therefore retained.[2] Following common practice in EMA studies [refs], all models were adjusted to account for linear drifts in affect ('Time in Study') and diurnal effects on affect ('Time of Day').[3–5]</td></tr>
<tr><td>Number</td><td>Pre-registered hypothesis</td><td>Pre-registered analysis plan</td><td>Deviation description</td></tr>
<tr><td>2a</td><td>That taking SSRIs modulates reward predictions by increasing the extent to which current mood biases said predictions.</td><td>Hypothesis 2a will be tested by constructing a separate linear mixed-effects model in order to assess the relationship between SSRI group membership and reward predictions, according to the specification described in formulae (9) and (10).<br><br>(9) [reward] prediction ~ SSRI + PA + SSRI:PA + (1+SSRI+PA+SSRI:PA | participant)<br><br>(10) [reward] prediction ~ SSRI + NA + SSRI:NA + (1+SSRI+NA+SSRI:NA | participant)</td><td>PA and NA were included as simultaneous predictors in a single model, with independent SSRI interaction terms for each, to account for their covariance. This also reduces the number of statistical tests relative to the pre-registered plan. An additional analysis included lagged RPEs to examine whether prior reward signals predict subsequent reward perception.[1]</td></tr>
<tr><td>2b</td><td>That SSRIs modulate participants' mood response to reward by altering the modelled relationship between positive and negative affect and reward prediction error and outcomes.</td><td>Hypothesis 2b will be tested by additionally incorporating SSRI group membership into the best-fit model arising from the tests of hypothesis 1 (e.g. those described by formulae (11) and (12)).<br><br>(11) $PA_{post}$ ~ $PA_{pre}$ + outcome + RPE + (outcome+RPE | participant/SSRI)<br><br>(12) $NA_{post}$ ~ $NA_{pre}$ + outcome + RPE + (outcome+RPE | participant/SSRI)</td><td>Unlike hypothesis 2a, PA and NA could not be combined into a single model given their roles as separate outcome variables here. To reduce the number of analyses, we focused on positive affect as the primary outcome, consistent with the theoretical emphasis on reward-driven mood under SSRI treatment.</td></tr>
<tr><td>2c</td><td>That the associations with SSRI usage are distinct for positive and negative prediction errors.</td><td>Hypothesis 2c will be tested by splitting the RPE term into distinct positive and negative RPE terms (e.g. formulae (13) and (14)).<br><br>(13) $PA_{post}$ ~ $PA_{pre}$ + outcome + $RPE_{pos}$ + $RPE_{neg}$ + (outcome+$RPE_{pos}$+$RPE_{neg}$ | participant/SSRI)<br><br>(14) $NA_{post}$ ~ $NA_{pre}$ + outcome + $RPE_{pos}$ + $RPE_{neg}$ + (outcome+$RPE_{pos}$+$RPE_{neg}$ | participant/SSRI)</td><td>As for 2b, analyses focus on positive affect to reduce the number of tests; the negative affect model was not estimated.</td></tr>
</table>

### Supplementary Information

**Supplementary Table 6. Controlling for unblinding.** Participants were able to correctly identify their group allocation at above-chance levels ($\chi^2 = 20.4$, $p < 0.001$).

| | Allocated placebo (n=30) | Allocated SSRI (n=40) |
|---|---|---|
| **Guessed placebo** | 19 (63%) | 9 (23%) |
| **Guessed SSRI** | 4 (13%) | 27 (67%) |
| **Uncertain** | 7 (24%) | 4 (10%) |
| **Total at follow-up (day 8)** | 27 (90%) | 36 (90%) |

### Supplementary Information

**Supplementary Table 7. Exploratory factor analysis of Mood Zoom** (two-factor solution; n = 3274).

| Item | Factor 1 | Factor 2 |
|---|---|---|
| Anxious | **0.51** | -0.01 |
| Happy | -0.02 | **0.87** |
| Irritable | **0.71** | -0.03 |
| Sad | **0.62** | -0.16 |
| Angry | **0.79** | 0.09 |
| Energetic | 0.03 | **0.60** |
| **Interpretation** | Negative affect | Positive affect |

***Note: Bold*** *indicates the loadings which dominate each factor (>0.4).*

### Supplementary Information

##### Pleasure and Mastery

**Supplementary Table 8. Exploratory factor analysis of EMA reward items** (two-factor solution; n = 3560).

| Item | Factor 1 | Factor 2 |
|---|---|---|
| Enjoyment | -0.08 | **0.77** |
| Social connectedness | 0.21 | **0.50** |
| Achievement | **0.67** | 0.27 |
| Effort | **0.80** | -0.17 |
| **Interpretation** | Mastery | Pleasure |

***Note: Bold*** *indicates the loadings which dominate each factor (>0.4).*

Pleasure and mastery prediction errors (PE) were computed for the corresponding reward dimensions. calculated as the difference between the actual experience of reward (r) and the expected reward (ν) extracted from paired post- and pre-activity surveys, respectively.

$$pleasure\ PE = pleasure\ \mathrm{r} - pleasure\ \mathrm{\nu}$$

$$mastery\ PE = mastery\ \mathrm{r} - mastery\ \mathrm{\nu}$$

To explore whether SSRIs act on distinct facets of reward to improve mood, the 'post-activity positive affect' model was rerun for pleasure and mastery reward dimensions. Specifically, post-activity positive affect was modelled as a function of pleasure $PE_t$, pleasure $r_t$, mastery $PE_t$, mastery $r_t$ and $PA_{t-1}$.

$$PA_t \sim (pleasure\ PE_t + pleasure\ \mathrm{r}_t + mastery\ PE_t + mastery\ \mathrm{r}_t + PA_{t-1}) * Group * Time\ in\ Study_t + Time\ of\ Day_t + Time\ in\ Study_t + (1 \mid participant)$$

#### Supplementary Information

**Supplementary Table 9. Estimated effects of SSRI, Pleasure and Mastery Outcome, Pleasure and Mastery Prediction Error, and Time in Study on positive affect**

$$PA_t \sim (pleasure\ PE_t + pleasure\ r_t + mastery\ PE_t + mastery\ r_t + PA_{t-1}) * Group * Time\ in\ Study_t + Time\ of\ Day_t + Time\ in\ Study_t + (1 \mid participant)$$

| Parameter | Effect | Standard Error | *t* | Degrees of freedom | p-value |
|---|---|---|---|---|---|
| Intercept | -0.03 | 0.08 | -0.31 | 55 | 0.756 |
| $PA_{t-1}$ | **0.30** | 0.04 | 8.48 | 1236 | **<0.001** |
| Pleasure $r_t$ | **0.22** | 0.03 | 6.30 | 1300 | **<0.001** |
| Mastery $r_t$ | **0.09** | 0.09 | 2.66 | 1304 | **0.008** |
| Pleasure $PE_t$ | 0.04 | 0.03 | 1.25 | 1272 | 0.213 |
| Mastery $PE_t$ | 0.02 | 0.03 | 0.53 | 1279 | 0.599 |
| SSRI | 0.04 | 0.10 | 0.41 | 55 | 0.686 |
| Time of $Day_t$ | **0.06** | 0.02 | 3.66 | 1253 | **<0.001** |
| Time in $Study_t$ | **-0.06** | 0.03 | -2.13 | 1276 | **0.033** |
| $PA_{t-1}$ * SSRI | -0.002 | 0.05 | -0.04 | 1261 | 0.967 |
| Pleasure $r_t$ * SSRI | **0.12** | 0.04 | 2.56 | 1297 | **0.010** |
| Mastery $r_t$ * SSRI | 0.002 | 0.04 | 0.04 | 1305 | 0.971 |
| Pleasure $PE_t$ * SSRI | 0.02 | 0.04 | 0.40 | 1273 | 0.695 |
| Mastery $PE_t$ * SSRI | -0.04 | 0.04 | -1.13 | 1278 | 0.259 |
| Time in $Study_t$ * SSRI | **0.07** | 0.04 | 2.06 | 1277 | **0.040** |
| Time in $Study_t$ * $PA_{t-1}$ | 0.03 | 0.03 | 1.14 | 1288 | 0.253 |
| Time in $Study_t$ * pleasure $r_t$ | -0.04 | 0.03 | -1.35 | 1263 | 0.179 |
| Time in $Study_t$ * mastery $r_t$ | -0.02 | 0.03 | -0.55 | 1269 | 0.583 |
| Time in $Study_t$ * pleasure $PE_t$ | -0.04 | 0.03 | -1.30 | 1258 | 0.197 |
| Time in $Study_t$ * mastery $PE_t$ | 0.04 | 0.03 | 1.47 | 1267 | 0.141 |
| $PA_{t-1}$ * SSRI * Time in $Study_t$ | 0.06 | 0.04 | 1.60 | 1283 | 0.109 |
| Pleasure $r_t$ * SSRI * Time in $Study_t$ | 0.004 | 0.04 | 0.09 | 1262 | 0.925 |
| Mastery $r_t$ * SSRI * Time in $Study_t$ | -0.001 | 0.04 | -0.01 | 1268 | 0.990 |
| Pleasure $PE_t$ * SSRI * Time in $Study_t$ | **0.10** | 0.04 | 2.63 | 1254 | **0.009** |
| Mastery $PE_t$ * SSRI * Time in $Study_t$ | -0.02 | 0.04 | -0.46 | 1266 | 0.644 |

***Note:*** *$PA_{t-1}$ = pre-activity positive affect; pleasure $r_t$ = post-activity pleasure outcome; pleasure $PE_t$ = post-activity pleasure Prediction Error; mastery $r_t$ = post-activity mastery outcome; mastery $PE_t$ = post-activity mastery Prediction Error; * = interaction;* ***bold*** *indicates significance at $p<0.05$.*

Mirroring the effects observed for enjoyment-related reward (Table 2), citalopram significantly increased the impact of pleasure on positive affect (pleasure $r_t$*SSRI: $\beta = 0.12$, $p < 0{,}010$), while also strengthening the impact of pleasure PEs on positive mood over time (pleasure $PE_t$*SSRI*Time in $Study_t$: $\beta = 0.10$, $p < 0.009$). In contrast, the only effect observed for mastery-related reward was a general increase in positive affect following positive mastery outcomes (mastery $r_t$: $\beta = 0.09$, $p < 0.008$). Together, these findings indicate that SSRIs may act on different facets of reward to improve momentary mood, primarily amplifying pleasure and enjoyment-related experiences, rather than mastery.

#### Supplementary Information

Notably, these findings echo those from a smartphone CBT study, which asked depressed patients to give ratings of pleasure and mastery when planning and evaluating a range of daily activities. Throughout the study, the authors found that level of pleasure that the patient expected was the most significant predictor of depression improvement.[6] These findings imply that behavioral interventions, and possibly SSRIs, may be most effective when targeting activities that generate inherent pleasure (e.g., going to a coffee shop), rather than effortful, achievement-focused tasks (e.g., tidying up the house). More broadly, these results highlight the utility in assessing reward as a multidimensional construct, given the dissociable effects of serotonin on pleasure- versus mastery-based reward processing, and the potential for tailored therapeutic interventions.
